# BATF2 is a regulator of interferon-γ signaling in astrocytes during neuroinflammation

**DOI:** 10.1101/2024.07.10.602938

**Authors:** Rachel A. Tinkey, Brandon C. Smith, Maria L. Habean, Jessica L. Williams

## Abstract

Astrocytic interferon (IFN)γ signaling is associated with a reduction in neuroinflammation. We have previously shown that the benefits of astrocytic IFNγ arise from a variety of mechanisms; however, downstream effectors responsible for regulating this protection are unknown. We address this by identifying a specific transcription factor that may play a key role in modulating the consequences of IFNγ signaling. RNA-sequencing of primary human astrocytes treated with IFNγ revealed basic leucine zipper ATF-like transcription factor (*BATF*)2 as a highly expressed interferon-specific gene. Primarily studied in the periphery, BATF2 has been shown to exert both inflammatory and protective functions; however, its function in the central nervous system (CNS) is unknown. Here, we demonstrate that human spinal cord astrocytes upregulate BATF2 transcript and protein in an IFNγ-specific manner. Additionally, we found that BATF2 prevents overexpression of interferon regulatory factor (IRF)1 and IRF1 targets such as Caspase-1, which are known downstream pro-inflammatory mediators. We also show that *Batf2*^−/−^ mice exhibit exacerbated clinical disease severity in a murine model of CNS autoimmunity, characterized by an increase in both CNS immune cell infiltration and demyelination. *Batf2*^−/−^ mice also exhibit increased astrocyte-specific expression of IRF1 and Caspase-1, suggesting an amplified interferon response *in vivo*. Further, we demonstrate that BATF2 is expressed primarily in astrocytes in MS lesions and that this expression is co-localized with IRF1. Collectively, our results further support a protective role for IFNγ and implicate BATF2 as a key suppressor of overactive immune signaling in astrocytes during neuroinflammation.

## Introduction

Astrocytes are the most abundant glial cell in the central nervous system (CNS) and are essential for both homeostatic function and response to injury and disease. Assuming diverse phenotypes, morphologies, and functional properties, astrocytes are uniquely poised to respond to complex cues in the CNS microenvironment that result in both protective and pathogenic outcomes^1^. Widespread astrocyte reactivity is a common pathogenic hallmark across a variety of neuroinflammatory diseases such as multiple sclerosis (MS), in which prolonged astrocyte activation contributes to early lesion formation and sustained inflammation^2,3^. While changes in astrocytic function can be stimulated by a variety of factors, a major contributor, particularly in neuroinflammatory disorders like MS, includes cytokines such as interferon (IFN)γ that are highly pleiotropic in nature. Astrocyte exposure to IFNγ has previously been shown to contribute to sustained inflammation and neuronal damage through initiation of cytokine production early in disease development^3–5^. However, IFNγ signaling in astrocytes has also been associated with protection from demyelination and clinical decline in progressive stages of immune-mediated neuroinflammation^6–8^. Further, we and others have recently shown additional protective roles for astrocytic IFNγ signaling, specifically in the spinal cord in an animal model of MS, experimental autoimmune encephalomyelitis (EAE)^9–12^.

While several avenues of astrocytic IFNγ signaling have been described, the molecules responsible for directly regulating this protection are currently unknown. A variety of downstream effectors, namely transcription factors, play a major role in the regulation of gene expression and can be directly stimulated by IFNγ. Using RNA-sequencing of primary human astrocytes treated with IFNγ, we identified the basic leucine zipper ATF-like transcription factor (*BATF*)2 as one of the most highly upregulated genes. BATF2 is a member of the basic leucine zipper (bZIP) family of transcription factors and was initially identified as an inhibitor of activator protein (AP)-1 function through direct interaction with c-JUN^13^. A multi-phenotypic immunomodulatory role for BATF2 has been demonstrated in peripheral models of inflammation in which BATF2 exerts both pro-inflammatory and protective functions. Specifically, BATF2 dimerization with c-JUN resulted in a reduction in interleukin (IL)-23 signaling during *Tryanosoma cruzi* infection^14^ and attenuated colon inflammation through promoting the degradation of signal transducer and activator of transcription (STAT)1 in mice affected with colitis^15^. Conversely, BATF2 has also been shown to interact with other IFNγ-inducible transcription factors, including interferon regulatory factor (IRF)1, which is known to promote macrophage activation and enhance inflammation in several contexts^16–18^.

Due to its known activity in response to cytokine stimulation and extensive regulatory role in the periphery, BATF2 is a prime candidate for modulation of neuroinflammatory events; however, the role of BATF2 in the CNS has not yet been described. In this study, we demonstrate that primary human astrocytes upregulate BATF2 in an IFNγ-specific manner and that IFNγ promotes nuclear expression of BATF2 which may regulate AP-1 transcription sites. This nuclear BATF2 then suppresses overexpression of IRF1 and IRF1 target genes, such as Caspase-1, that have been implicated as pro-inflammatory and deleterious during autoimmune demyelination^19–23^. Utilizing EAE as a model for neuroinflammation, we also show that *Batf2*^−/−^ mice exhibit exacerbated EAE severity, characterized by an increase in both peripheral immune cell infiltration and demyelination compared to *Batf2*^+/+^ controls. Further, *Batf2*^−/−^ mice also exhibit increased astrocyte-specific expression of both IRF1 and Caspase-1, suggesting an increased inflammatory phenotype driven, in part, by an enhanced interferon response *in vivo*. BATF2 was also found to be primarily expressed by astrocytes in MS lesions and colocalize with IRF1. Taken together, these data suggest that IFNγ-mediated BATF2 expression in astrocytes may directly facilitate a neuroprotective microenvironment downstream of IFNγ signaling during MS and EAE. These findings further enhance our current understanding of immune-mediated CNS repair mechanisms driven by alternate astrocytic states during neuroinflammatory disease.

## Results

### IFNγ enhances BATF2 expression in human astrocytes

We have previously identified protective mechanisms of IFNγ in astrocytes during chronic autoimmunity^9,24^; however, the specific intrinsic mechanisms responsible for regulating these pathways have not been fully elucidated. To investigate potential regulators of astrocytic IFNγ signaling, we performed RNA sequencing of primary human astrocytes (**Supplemental Figure 1A**) treated with cytokines known to be present during a variety of neuroinflammatory events including IFNγ, tumor necrosis factor (TNF)α, IL-1β, IL-17, and granulocyte-macrophage colony-stimulating factor (GM-CSF). Since the primary pathological target during EAE is the spinal cord and over 70% of MS patients present with spinal cord lesions^25,26^, we focused our analysis on this CNS region. Principal component analysis (PCA) revealed a heterogenous response of spinal cord astrocytes to different cytokines with IFNγ treatment having the greatest separation from media, followed by TNFα and IL-1β, while IL-17 and GM-CSF remained the most similar to control-treated astrocytes (**Figure 1A**). To highlight genes specific to IFNγ, the top upregulated genes were analyzed compared to media-treated and plotted on a heat map comparing expression patterns to TNFα and IL-1β (**Figure 1B**). Genes exclusive to IFNγ treatment included *HLA-DMB*, *HLA-DQB1, HLA-DOA, INHBE, IL18BP,* and *BATF2* (**Figure 1C**). *HLA-DMB*, *HLA-DQB1,* and *HLA-DOA* transcribe proteins involved in MHC class II antigen presentation^27^, while *INHBE* transcribes the preproprotein inhibin subunit beta E, which is primarily expressed in the pancreas and liver and is a member of the transforming growth factor β superfamily^28^. Further, the *IL18BP* gene transcribes IL-18 binding protein that functions to inhibit IL-18 signaling^29^. Interestingly, of the genes identified as being IFNγ-specific, *BATF2* was the only gene responsible for producing a transcription factor and BATF2 had previously been shown to regulate inflammatory processes in the periphery^14,16,17,30^. To confirm the transcriptional regulation of BATF2 by IFNγ, we treated primary human astrocytes over time and found that peak *BATF2* expression occurred at 12-24h post-treatment (**Figure 1D**). Importantly, *BATF2* expression was specific to IFNγ compared to other inflammatory mediators and BATF family members (**Figure 1E**). While a dose titration using types I, II, and III interferons revealed that *BATF2* transcript was upregulated in astrocytes by both IFNγ and IFNβ, IFNγ was significantly more proficient at promoting *BATF2* expression (**Supplemental Figure 1B**). Further, astrocyte BATF2 protein levels were elevated 24h following IFNγ stimulation (**Figure 1F-G**). These data suggest that IFNγ signaling in astrocytes is sufficient to upregulate the transcription and protein expression of BATF2 in a specific manner.

**Figure 1.**
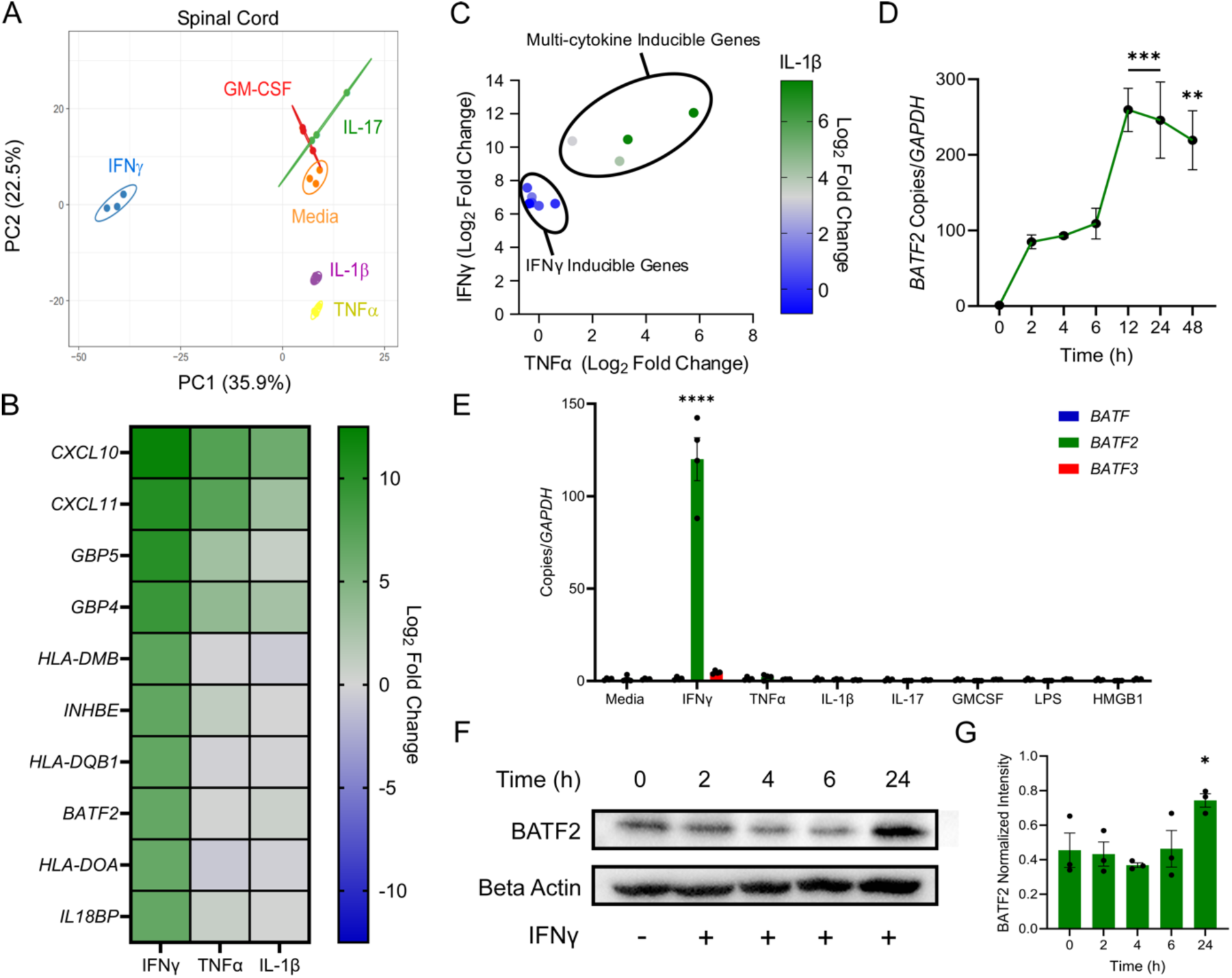
IFNγ signaling induces BATF2 expression in astrocytes. (A) PCA analysis of RNA sequencing demonstrating distinct gene expression patterns of human spinal cord astrocytes treated with 10ng/mL IFNγ, TNFα, IL-1β, IL-17, GM-CSF, and media for 24h. (B) Heat map of top upregulated genes in human spinal cord astrocytes treated with 10ng/mL of IFNγ for 24h compared to TNFα and IL-1β treatments. Data represent Log2 fold change compared to media from 3 independent samples. (C) Top upregulated genes in human spinal cord astrocytes treated with 10ng/mL IFNγ for 24h compared to TNFα and IL-1β treatments. Data represent the Log2 fold change compared to media from 3 independent samples. (D) *BATF2* transcript levels of human spinal cord astrocytes stimulated with 10ng/mL IFNγ for 0-48h. \*\**p* < 0.01 and \*\*\**p* < 0.001 compared to media-treated samples by one-way ANOVA. Data points represent mean ± SEM (*n*=3). (E) *BATF2* transcript levels of human spinal cord astrocytes treated with 10ng/mL of various inflammatory stimuli for 24h. \*\*\*\**p* < 0.0001 compared to media-treated samples by two-way ANOVA. Bars represent mean ± SEM (*n*=4). (F) Representative western blot of whole cell BATF2 protein levels in human spinal cord astrocytes treated with 10ng/mL of IFNγ for 24h. (G) Quantification of whole cell BATF2 protein levels in primary human spinal cord astrocytes shown in F, normalized to β-actin expression. \**p* < 0.05 compared to media-treated samples by one-way ANOVA. Bars represent mean ± SEM (*n*=3).

### IFNγ regulates BATF2 transcriptional activity

The function of BATF2 in the CNS and in astrocytes has yet to be explored. To determine the chromatin occupancy of BATF2 at specific gene targets downstream of IFNγ, we treated primary human spinal cord astrocytes with IFNγ or a media control for 24h and processed samples for chromatin immunoprecipitation (ChIP) sequencing, which was performed and analyzed by Active Motif. Peak regions for each treatment were determined using MACS2 software and mapped to the hg18 human genome. Correlation coefficients of peaks between treatments demonstrated high congruency between media and IFNγ samples (**Supplemental Figure 2A**), indicating high reproducibility among replicates. A relatively high correlation between media and IFNγ samples was also found, but not unexpected, as BATF2 is known to be constitutively expressed, suggesting a potential overlap in transcriptional targets between media and IFNγ treatments. To tease out specific differences in peak metrics between media and IFNγ samples, intervals of genomic regions with local enrichments in BATF2 tag numbers were grouped into merged peak regions. Notably, human astrocytes treated with IFNγ demonstrated an increased number of merged peak regions compared to media controls (**Figure 2A**). Specifically, 326 merged regions were found to be IFNγ-specific, compared to 2 regions within media samples, and 97 regions were overlapped across all replicates of each treatment group (**Supplemental Figure 2B**). Additionally, IFNγ-treated astrocytes also demonstrated a higher average peak tag number per merged region (**Figure 2B**) suggesting increased BATF2 chromatin accessibility and transcriptional activity with IFNγ stimulation. Peak enrichments at specific genomic loci were also identified and compared between media– and IFNγ-treated samples. Human astrocytes treated with IFNγ had an overall increase in BATF2 binding events throughout the genome as well as specific enrichments at both promoter regions and CpGI islands (**Figure 2C**). This suggests that there is increased recruitment of BATF2 to upstream and downstream transcriptional sites that have a direct impact on gene expression. Further, DNA motif analysis of media-stimulated astrocytes showed BATF2 binding at nuclear factor (NF)Y sites associated with CCAAT boxes (**Figure 2D**). NFY is a highly conserved heterotrimeric transcription factor necessary for the expression of genes involved in the cell cycle, DNA repair, and transcription initiation^31–33^. Conversely, following IFNγ treatment, we observed a significant increase in BATF2 binding at AP-1 sites (**Figure 2E**), suggesting that IFNγ-mediated BATF2 regulation of AP-1 may influence proliferation and/or critical inflammatory pathways in the CNS^34–37^. Taken together, these data suggest that IFNγ induces the transcriptional activity of BATF2 and shifts BATF2 function from modulation of normal cellular processes to those involved in neuroinflammation.

**Figure 2.**
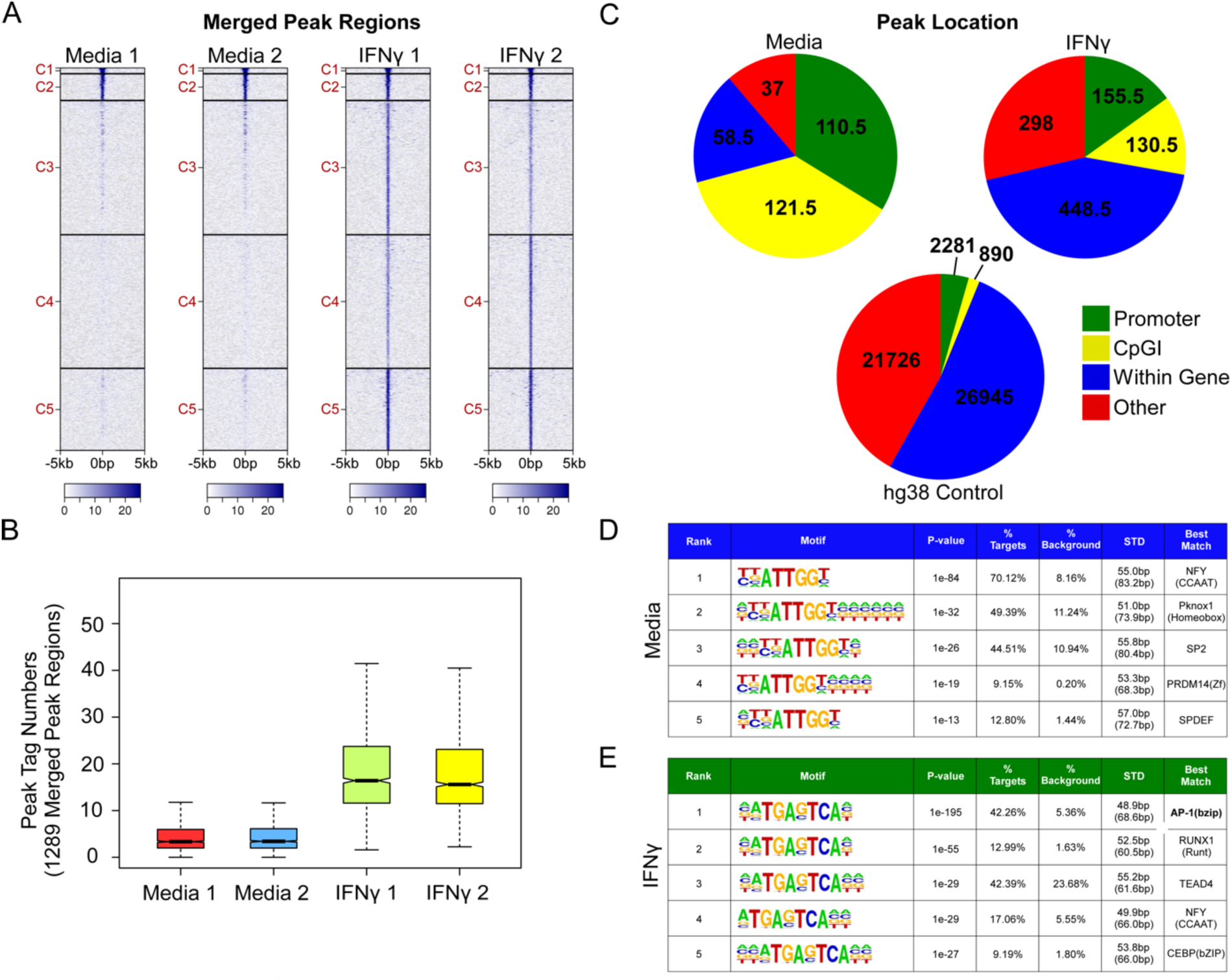
IFNγ increases DNA binding of BATF2 at inflammatory sites in astrocytes. (A) Merged peak regions heat map of BATF2 binding in human spinal cord astrocytes treated with 10ng/mL IFNγ or media for 24h. Scale bar indicates BATF2 binding events per merged region. (B) Peak tag numbers of BATF2 binding events in 1289 merged peak regions of human spinal cord astrocytes treated with 10ng/mL IFNγ or media for 24h. (C) Peak location of BATF2 binding events in human spinal cord astrocytes treated with 10ng/mL IFNγ or media for 24h. hg38 binding events were used as an internal control. Data are the average peak locations between two independent replicates per treatment. (D-E) Top BATF2 binding motifs of media– and IFNγ-treated human spinal cord astrocytes. Data shown in A-E are representative of two independent samples per treatment.

### BATF2 suppresses IRF1 over-expression downstream of IFNγ in astrocytes

To identify specific pathways regulated by BATF2 downstream of IFNγ, we performed ingenuity pathway analysis (IPA) on annotated genes bound upstream or downstream by BATF2 in treated astrocytes. IPA identified several inflammatory pathways significantly associated with BATF2 bound regions including the IFNγ and MS signaling pathways (**Figure 3A**). Additionally, regions associated with BATF2 binding were predicted to be activated by upstream regulators such as IFNγ and have both direct and indirect interactions with a variety of other immune function modulators (**Figure 3B**). This may suggest a potential role for BATF2 in several inflammatory mechanisms adjacent to IFNγ signaling. Further analysis of the IFNγ and MS pathways (**Figure 3C-D**) identified IRF1 as a common gene associated with increased BATF2 binding at upstream CpGI island regions in astrocytes stimulated with IFNγ compared to media controls (**Figure 3E**). To validate how the IRF1 signaling axis was regulated by BATF2, control or *Batf2*^−/−^ murine astrocytes were stimulated with media or IFNγ for 48h and levels of *Irf1* transcript, as well as IRF1 target genes including *Psmb8*, *Casp1*, *Mhc1*, *Tap1*, and *Tap2* were quantified. Transcript levels of *Batf2* were also quantified as a control (**Supplemental Figure 3A**). Notably, loss of BATF2 resulted in increased expression of IRF1 as well as downstream IRF1 targets (**Figure 3F-K**). These data suggest that BATF2 is necessary to keep IFNγ-inducible genes such as IRF1 in check and that deletion of BATF2 results in amplified IFNγ signaling in astrocytes.

**Figure 3.**
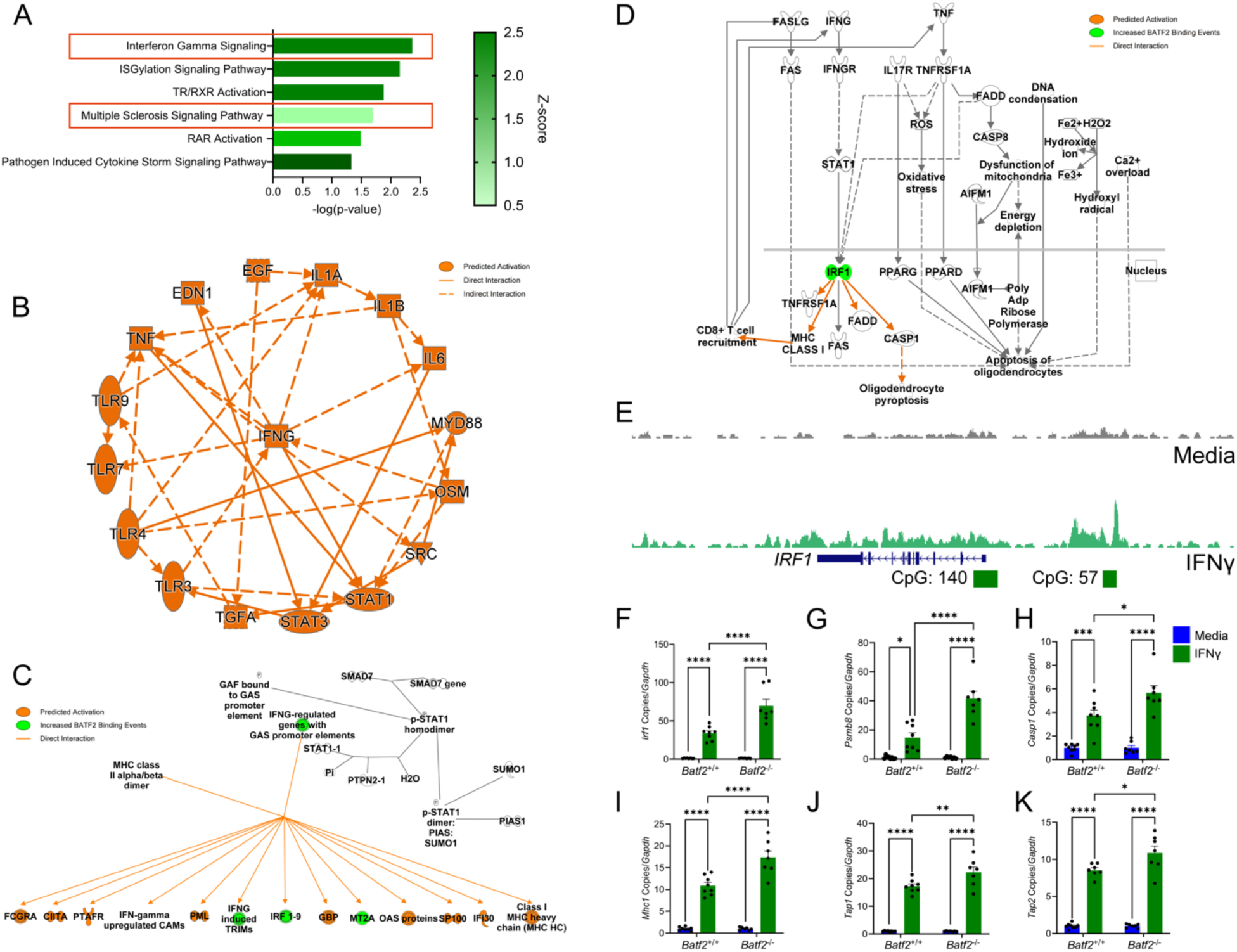
Astrocytic BATF2 suppresses the IRF1 signaling axis downstream of IFNγ. (A) Ingenuity pathway analysis of top regulated pathways by BATF2 in human spinal cord astrocytes stimulated with 10ng/mL IFNγ for 24h. (B) Graphical summary of predicated targets regulated by BATF2 in human spinal cord astrocytes stimulated with 10ng/mL IFNγ for 24h. (C) Adapted Ingenuity Pathway Analysis: Interferon Gamma Signaling pathway from panel A. (D) Adapted Ingenuity Pathway Analysis: Multiple Sclerosis Signaling Pathway from panel A. (E) Peak region counts of BATF2 binding events upstream of the *IRF1* gene locus and at CpG islands in human spinal cord astrocytes stimulated with 10ng/mL IFNγ for 24h (green) or media (grey). (F-K) Quantification of *Irf1* and target gene transcript expression in *Batf2*^+/+^ and *Batf2*^−/−^ primary murine spinal cord astrocytes stimulated with media or 10ng/mL IFNγ for 48h. Data is representative of 2 independent experiments. Individual data points were normalized to the media average and are representative of individual mice. \**p* < 0.05, \*\**p* < 0.01, \*\*\**p* < 0.001, ****P < 0.0001 compared to media-treated samples by two-way ANOVA. Bars represent mean ± SEM (*n*=7). Data shown in A-E are representative of two independent samples per treatment.

### Deletion of BATF2 exacerbates chronic EAE

We and others have previously shown that dysregulated IFNγ signaling leads to exacerbation of EAE^7–9,24^. Moreover, with a focus on peripheral immune cells, other BATF family members have been studied in the context of EAE^38,39^. However, the impact of BATF2 on the CNS during neuroinflammation has yet to be described. Initially, we wanted to confirm that BATF2 expression was dependent on IFNγ signaling in astrocytes *in vivo*. To validate this, we injected TdTomato::*Aldh1l1*-Cre^ERT2+^ *Ifngr1*^fl/fl^ mice^9^ and littermate controls with tamoxifen to conditionally delete the IFNγ receptor on astrocytes. Following administration of tamoxifen, mice were immunized for EAE and BATF2 expression was assessed. As expected, TdTomato::*Aldh1l1*-Cre^ERT2+^ *Ifngr1*^fl/fl^ mice exhibited exacerbated EAE severity compared to littermate controls (**Supplemental Figure 4A**). Importantly, there was a significant decrease in both total BATF2 expression and BATF2 colocalized with TdTomato^+^ astrocytes within lesions of *Ifngr1*^fl/fl^ *Aldh1l1*-Cre^ERT2+^ mice compared to littermate controls (**Supplemental Figure 4B-E**). This suggests that full BATF2 expression in astrocytes requires IFNγ. To further address how BATF2 directly modulates neuroinflammation, we induced EAE in *Batf2*^−/−^ mice (**Supplemental Figure 3B-C**) and controls. Following EAE induction, *Batf2*^−/−^ mice exhibited an exacerbated clinical disease course (**Figure 4A-B**), as well as increased demyelination, lesion size, and immune cell infiltration compared to littermate controls (**Figure 4C-I**). To assess if *Batf2*^−/−^ mice had dysregulated IFNγ signaling, we quantified IRF1 and Caspase-1 protein levels in the ventral spinal cord white matter tracks and found that *Batf2*^−/−^ mice had a significant increase in both IRF1 and Caspase-1 expression within EAE lesions. Importantly, this increase was observed specifically in astrocytes, validating previous *in vitro* findings (**Figure 4J-Q**). Taken together, these data suggest that BATF2 is protective during neuroinflammation and may act as a negative regulator of IFNγ-induced inflammatory pathways *in vivo* to limit immune-mediated CNS injury.

**Figure 4.**
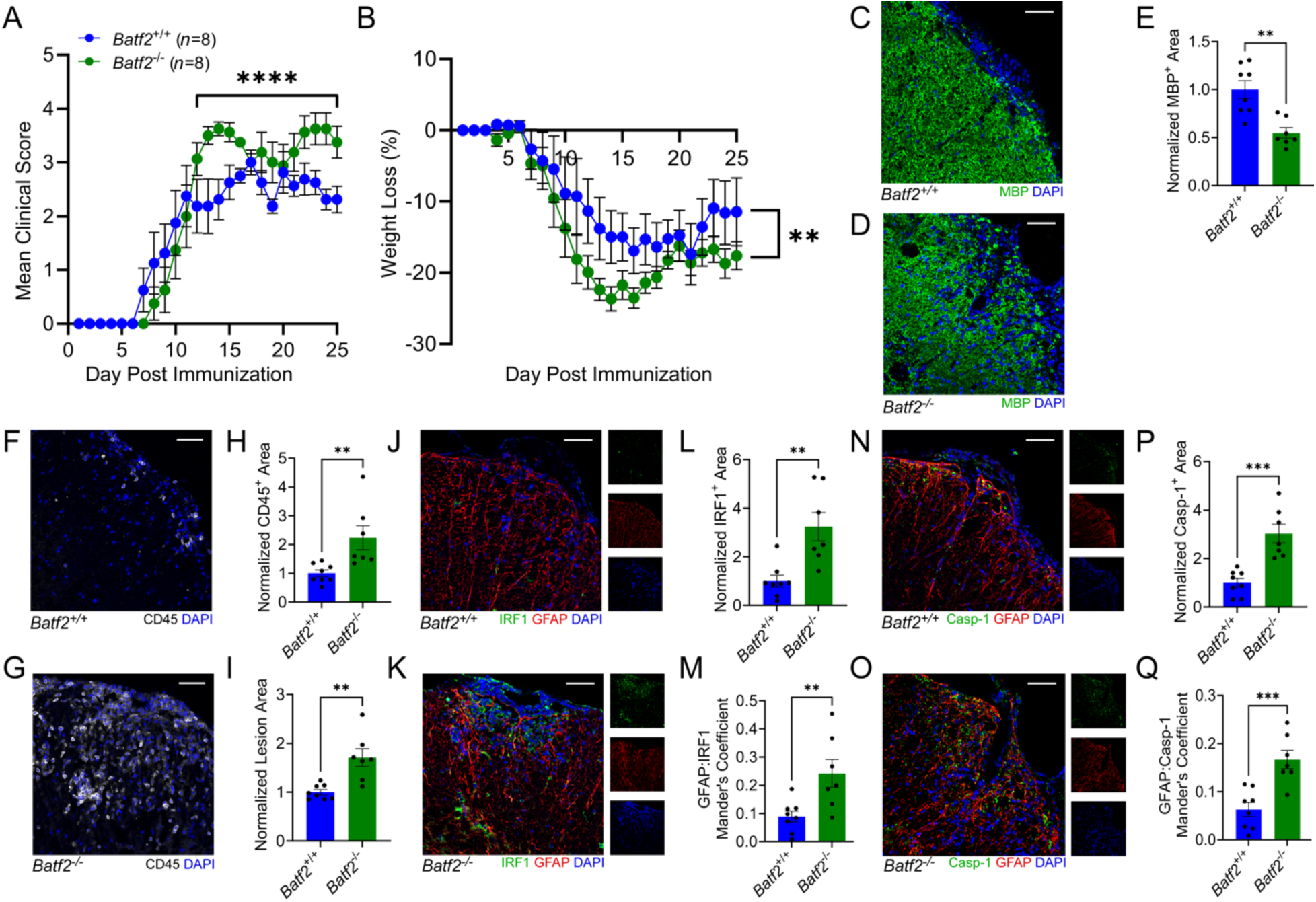
Loss of BATF2 exacerbates EAE and enhances IRF1 and Caspase-1 expression in vivo. (A-B) EAE was induced in *Batf2*^−/−^ and control mice. EAE clinical course (A) and weights (B) were blindly monitored. Data are a combination of 3 independent experiments and were analyzed using the Mann–Whitney *U* test for nonparametric data. \*\**p* < 0.01 from day 2, \*\*\*\**p* < 0.0001. Data points represent mean ± SEM (*n*=8 per genotype). 25 days post-immunization, mice were sacrificed and the CNS tissue was cryopreserved for immunofluorescent analysis. Ventral white matter tracts of the lumbar spinal cord were imaged using confocal microscopy. (C-D) Spinal cord tissue from (C) *Batf2*^+/+^ and (D) *Batf2*^−/−^ mice labeled for MBP and nuclei counterstained with DAPI. Scale bars, 50μm. (E) Quantification of MBP area normalized to the average of *Batf2*^+/+^ control mice. \*\**p* < 0.01 compared to *Batf2*^+/+^ samples by two-tailed Student’s *t* test. Bars represent mean ± SEM. (F-G) Spinal cord tissue from (F) *Batf2*^+/+^ and (G) *Batf2*^−/−^ mice labeled for CD45 and nuclei counterstained with DAPI. Scale bars, 50μm. (H-I) Quantification of CD45 area and lesion area normalized to the average of *Batf2*^+/+^ control mice. \*\**p* < 0.01 compared to *Batf2*^+/+^ samples by two-tailed Student’s *t* test. Bars represent mean ± SEM. (J-K) Spinal cord tissue from (J) *Batf2*^+/+^ and (K) *Batf2*^−/−^ mice labeled for IRF1, GFAP, and nuclei counterstained with DAPI. Scale bars, 50μm. (L-M) Quantification of IRF1 area normalized to the average of *Batf2*^+/+^ control mice and colocalization of GFAP and IRF1 for *Batf2*^+/+^ and *Batf2*^−/−^ mice. \*\**p* < 0.01 compared to *Batf2*^+/+^ samples by two-tailed Student’s *t* test. Bars represent mean ± SEM. (N-O) Spinal cord tissue from (N) *Batf2*^+/+^ and (O) *Batf2*^−/−^ mice labeled for Caspase-1, GFAP, and nuclei counterstained with DAPI. Scale bars, 50μm. (P-Q) Quantification of Caspase-1 area normalized to the average of *Batf2*^+/+^ control mice and colocalization of GFAP and Caspase-1 for *Batf2*^+/+^ and *Batf2*^−/−^ mice. ***P < 0.001 compared to *Batf2*^+/+^ samples by two-tailed Student’s *t* test. Bars represent mean ± SEM. Data in (C-Q) are representative of 3 independent experiments and include mice that survived until endpoint at day 25 (*n*=8 for *Batf2*^+/+^, *n*=7 for *Batf2*^−/−^). Each data point is representative of an individual mouse.

### BATF2 is expressed in astrocytes within MS lesions

To determine if BATF2 may also have a role in MS, we first characterized its expression in chronic active white matter lesions. These lesions are one of the hallmarks of the chronic, progressive stage of disease in which IFNγ signaling is thought to confer protection^6–9,24^. Using post-mortem MS tissue (**Table 1**) we identified chronic active lesions and normal appearing white matter (NAWM) based on the presence or lack of myelin basic protein (MBP) and the classic myeloid cell rim using ionized calcium binding adaptor molecule (IBA)1 labeling (**Figure 5A-B**). We then assessed the lesion rim, lesion core, and NAWM using immunofluorescence imaging of BATF2 (**Figure 5C-Ei**) and found that BATF2 was significantly upregulated in the lesion compared to NAWM. Within the lesion, BATF2 was more highly expressed in the core compared to the rim (**Figure 5F**). Colocalization analysis demonstrated that BATF2 expression within the core was more closely associated with ALDH1L1^+^ astrocytes compared to IBA1^+^ myeloid cells while expression in the rim was relatively similar among cell types (**Figure 5G**). Next, we assessed the expression pattern of IRF1 in relation to BATF2 (**Figure 5H-Ii**). Like BATF2, we found that IRF1 was more readily expressed in the lesion compared to NAWM and tended to be higher in the lesion core compared to the rim (**Figure 5H-K**). Further quantification again revealed that IRF1 localized to lesion core-associated Sox9^+^ astrocyte nuclei compared to IBA1^+^ myeloid cells (**Figure 5J,L**). Finally, regardless of sublesion location, we found that IRF1 colocalized with BATF2 (**Figure 5M**). These data demonstrate that both BATF2 and IRF1 are expressed in astrocytes in chronic active lesions and their co-expression may suggest that downstream of IFNγ signaling, BATF2 might regulate IRF1 expression during MS to prevent excessive inflammation, particularly in the lesion core.

**Figure 5.**
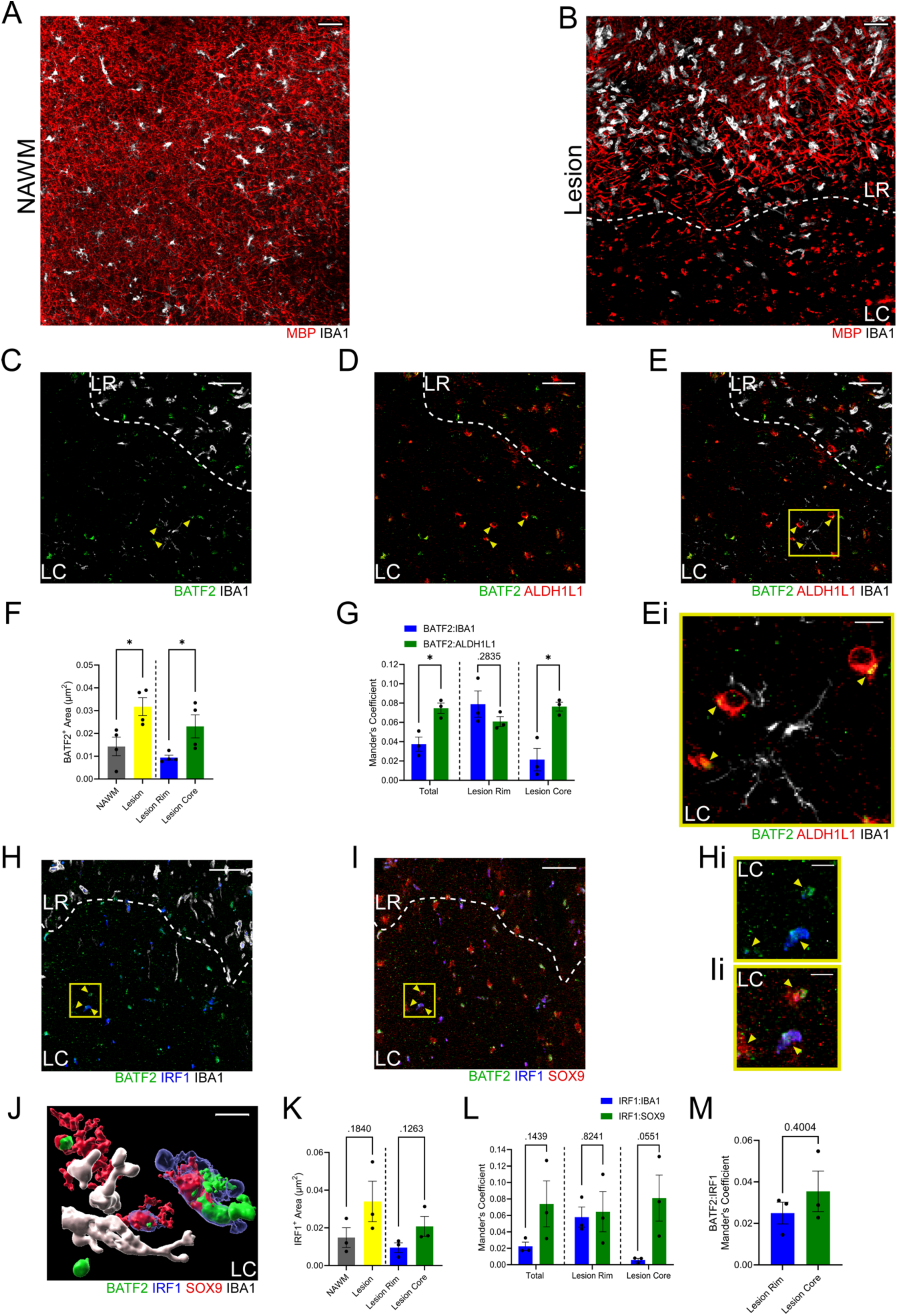
BATF2 is expressed by astrocytes in MS lesions and colocalizes with IRF1. (A-B) Human post-mortem MS brain sections (*n*=4) labeled for MBP and IBA1 to identify NAWM and chronic active lesions. LR=lesion rim, LC=lesion core. Scale bars, 50μm. (C-E) Chronic active lesions labeled for BATF2, IBA1, and ALDH1L1. LR=lesion rim, LC=lesion core. Scale bars, 50μm. (Ei) Digital zoom of a chronic active lesion labeled for BATF2, IBA1, and ALDH1L1. LC=lesion core. Scale bar, 10μm. (F) Quantification of BATF2 area. \**p* < 0.05 by two-tailed Student’s *t* test for each set of bars. Bars represent mean ± SEM. Data points are representative of all patients from Table 1. (G) Quantification of colocalization between BATF2 and ALDH1L1 (green) and BATF2 and IBA1 (blue). \**p* < 0.05 by two-tailed Student’s *t* test for each set of bars. Bars represent mean ± SEM. Data points are representative of patients MS52, MS115, and MS18 from Table 1. (H-I) Chronic active lesion labeled for BATF2, IRF1, IBA1, and Sox9. LR=lesion rim, LC=lesion core. Scale bars, 50μm. (Hi-Ii) Digital zoom of chronic active lesions labeled for BATF2, IRF1, IBA1, and Sox9. LC=lesion core. Scale bar, 10μm. (J) 3D render of a chronic active lesion labeled for BATF2, IRF1, IBA1, and Sox9. Scale bar, 10μm. (K) Quantification of IRF1 Area. Data was analyzed by a two-tailed Student’s *t* test for each set of bars. Bars represent mean ± SEM. Data points are representative of patients MS52, MS18, and MS160 from Table 1. (L) Quantification of colocalization between IRF1 and Sox9 (green) and IRF1 and IBA1 (blue). Data was analyzed by two-tailed Student’s *t* test for each set of bars. Bars represent mean ± SEM. Data points are representative of patients MS52, MS18, and MS160 from Table 1. (M) Quantification of colocalization of BATF2 and IRF1 in the lesion rim and lesion core. Data was analyzed by a two-tailed Student’s *t* test. Bars represent mean ± SEM. Data points are representative of patients MS52, MS18, and MS160 from Table 1.

**Table 1.**
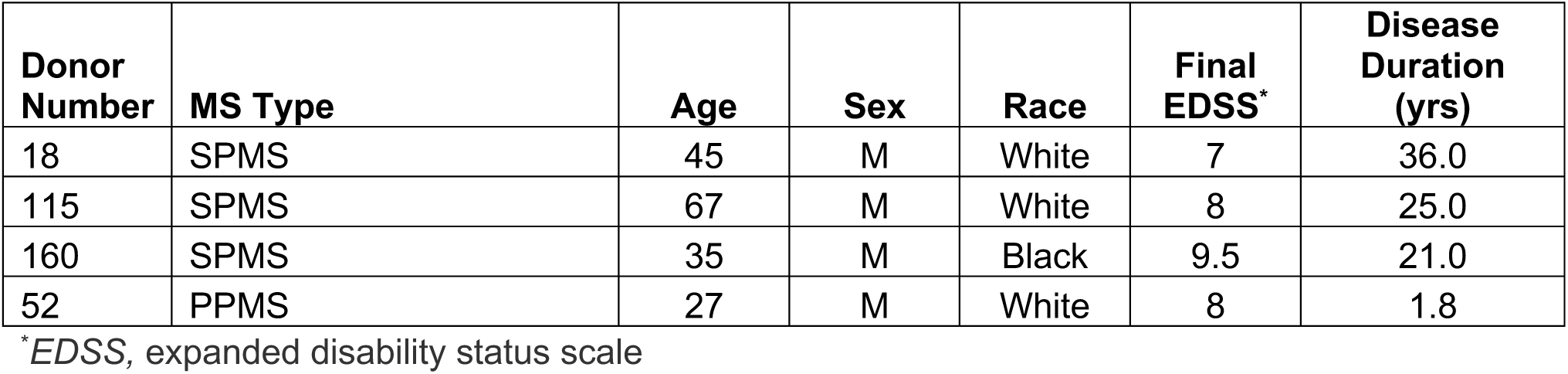
Patient Characteristics.

| **Donor Number** | **MS Type** | **Age** | **Sex** | **Race** | **Final EDSS\*** | **Disease Duration (yrs)** |
| --- | --- | --- | --- | --- | --- | --- |
| 18 | SPMS | 45 | M | White | 7 | 36.0 |
| 115 | SPMS | 67 | M | White | 8 | 25.0 |
| 160 | SPMS | 35 | M | Black | 9.5 | 21.0 |
| 52 | PPMS | 27 | M | White | 8 | 1.8 |

\*EDSS, expanded disability status scale

## Discussion and Conclusions

Growing evidence suggests that astrocytic IFNγ signaling is protective during chronic CNS autoimmunity^7–9,24^. In this study, we extend these findings to reveal a novel transcriptional regulator of IFNγ-mediated protection in astrocytes. Our study demonstrates that the transcription factor BATF2 is upregulated in response to IFNγ stimulation by astrocytes and prevents overabundant transcription of interferon-specific genes such as IRF1 and its downstream targets. Furthermore, loss of BATF2 led to exacerbated EAE with an increase in peripheral immune cell infiltration, demyelination, and expression of IRF1 and Caspase-1 by astrocytes within lesions. Colocalized BATF2 and IRF1 expression in astrocytes was also found to occur within chronic active MS lesions, suggesting a similar protective mechanism in human neuroinflammatory disease.

To date, the vast majority of BATF2 literature has been focused on its role in the periphery, to potentiate inflammatory responses in the context of infection. Of note, BATF2 has been shown to facilitate expression of inflammatory genes such as *Tnf*, *Ccl5*, and *Nos2* in bone marrow-derived macrophages downstream of IFNγ and during Mycobacterium tuberculosis (Mtb) infection *in vitro*. This effect was thought to result from BATF2 dimerization with IRF1, suggesting that a pro-inflammatory BATF2-IRF1 signaling axis exists in macrophages^16^. Further *in vivo* studies of Mtb infection in BATF2-deficient mice demonstrated reduced inflammatory chemokine and cytokine responses in lung macrophages, leading to decreased pulmonary inflammation and increased survival suggesting that a BATF2-associated inflammatory signature contributes to Mtb-associated mortality^18^. Contradictory to this evidence, during acute murine schistosomiasis infection, loss of BATF2 resulted in an exacerbated immune response leading to increased TNFα secretion by cells of the small intestine and enhanced T cell and dendritic cell activation^18^. A similar phenotype was observed in a murine model of colitis in which BATF2 deficiency increased CCL2 secretion through overactivation of STAT1 in intestinal epithelial cells^15^. Together, these previous studies suggest that BATF2 function is dependent on the context and nature of the immune response as it has been shown to promote inflammation downstream of Type 1 responses but is immunomodulatory during Type 2 immunity in the periphery. Our data add complexity to BATF2 function as we demonstrate a protective role for BATF2 in astrocytes downstream of IFNγ, a classic Type 1 neuroinflammatory response, in an animal model of MS contradicting peripheral phenotypes. We also show that BATF2 suppresses overactive IRF1 signaling in astrocytes rather than the synergistic pro-inflammatory BATF2-IRF1 interaction seen in macrophages. These discrepancies suggest that cells of the CNS respond uniquely to inflammatory stimuli and that the pathological location and immune response type may also influence whether BATF2 functions in a deleterious or protective capacity. Further, these functional disparities could be driven by cell type as astrocytes have been shown to respond differently to IFNα stimulation compared to myeloid cells such as microglia^40^. As such, it is possible that these distinctions extend to type II and III interferons, as well as other cytokine families, and may point to fine-tuned specialization in CNS cell function that contribute to the divergent BATF2 response observed across cell types.

Within the context of neuroinflammatory disease, our study highlights the importance of BATF2 in the regulation of active interferon signaling during CNS autoimmunity as we have shown that BATF2 negatively regulates critical inflammatory genes downstream of IFNγ in astrocytes. These genes include *Irf1* and *Casp1* which are known to perpetuate inflammatory cascades contributing to tissue damage during EAE^19–23^. Specifically, IRF1 and Caspase-1 colocalize in astrocytes within demyelinating lesions and mice lacking IRF1 failed to induce Caspase-1 expression and were resistant to EAE. This observation may also be facilitated by CNS resident cells as bone marrow chimera experiments revealed that only *Irf1*^−/−^ mice receiving wildtype bone marrow were protected from severe EAE compared to controls^20^. Further, expression of a dominant-negative form of IRF1 in oligodendrocytes ameliorated EAE and contributed to a reduced lesion load and parenchymal infiltration, confining immune cells largely to the perivascular space^21^. This suggests that IRF1 and subsequent activation of downstream genes may contribute to chemotaxis as well as parenchymal invasion of peripheral immune cells during neuroinflammation. Our findings corroborate these studies as we demonstrate that *Batf2*^−/−^ mice express increased IRF1 and Caspase-1 and have more extensive parenchymal infiltration during EAE compared to controls, implying that BATF2 suppression of IRF1 may work to limit CNS immune cell infiltration.

IRF1 single nucleotide polymorphisms are also associated with progressive MS^41^ and IRF1 expression has been characterized within glial cells in chronic active MS lesions^20^. However, little is known regarding how IRF1 expression is regulated at the cellular level within the lesion microenvironment. Our study demonstrated that IRF1 and BATF2 expression were frequently colocalized in glial cells within chronic active lesions. Interestingly, this colocalization occurred most frequently in the lesion core within astrocytes suggesting that BATF2 may be upregulated to suppress IRF1-induced inflammation in the quiescent lesion core. Cytokines including IFNγ are known inducers of astrocyte activation and may be released by activated myeloid cells within the lesion rim and contribute to scar formation^42,43^. While glial scarring is thought to inhibit remyelination, it is also essential for inflammatory confinement and protection of damaged axons^44–47^. Given the demonstrated protective capacity of astrocytic BATF2 downstream of IFNγ signaling and colocalization with inflammatory factors such as IRF1 in MS lesions, the specific upregulation of BATF2 within the lesion core may act as a facilitator of tissue sparing or repair.

Finally, loss of BATF2 has been linked with hyperactive interferon signaling and subsequent development of epilepsy and severe cognitive disability. Specifically, patients with a homozygous stop-gain mutation of *BATF2* presented with upregulated interferon-specific gene signatures in whole blood, resembling a type I interferon phenotype that is closely associated with interferonopathy^48^. Additionally, peripheral immune cells of patients with non-functional BATF2 were hyper-responsive to toll like receptor activation and overexpressed inflammatory cytokines including IFNα, TNF, IL-6 and IL-23^48^. This may indicate that BATF2 has the ability to cross regulate other inflammatory pathways in addition to those directly downstream of interferons. Interestingly, IFNγ can suppress IL-1β signaling through a variety of mechanisms including the inhibition of AP-1 and downstream mitogen activated protein kinase pathways^49–52^. Astrocytic BATF2 may therefore be a key regulator of other neuroinflammatory signaling mechanisms which may be applicable across many neurological disorders and warrants further investigation.

## Acknowledgements

We thank Kaitlin Kaiser for her assistance in mouse colony management, husbandry, genotyping, and contributions to the graphical abstract design. We also thank Dr. Bruce Trapp and Dr. Ranjan Dutta for sourcing of human MS tissue, as well as Orion Brock and Dr. Benjamin Shaw for insightful discussion. This work was supported by National Institute of Neurological Disorders and Stroke (NINDS) R01 NS119178 (awarded to Jessica L. Williams), R35 NS097303 (awarded to Bruce D. Trapp), and National MS Society RFA-2203-39228 (awarded to Jessica L. Williams).

## Author Contributions

RAT: Conceptualization, Data curation, Formal analysis, Writing – original draft, Writing – review & editing. BCS: Conceptualization and Data curation. MLH: Data curation. JLW: Conceptualization, Data curation, Funding acquisition, Writing – review & editing.

## Resource availability

### Lead contact

Further information and requests for resources and reagents should be directed to and will be fulfilled by the lead contact, Jessica L. Williams.

### Materials availability

All biological resources, antibodies, cell lines and model organisms and tools are either available through commercial sources or available upon request from the lead contact. Further information and requests for resources and reagents listed in Key Resources Table should be directed to the Lead Contact.

### Data and code availability

● The data sets generated during the current study are available in this paper or from the lead contact upon reasonable request.
● This paper does not report original code.
● Any additional information required to reanalyze the data reported here is available from the lead contact upon request.

## Experimental model and study participant details

### Animals

C57BL/6NJ and *Batf2*^−/−^ mice (strain no. 031903) were obtained from The Jackson Laboratories. All mice were subjected to EAE induction at 9-10 weeks of age. All housing, breeding, and procedures were approved by the Institutional Animal Care and Use Committee at the Lerner Research Institute, Cleveland Clinic Foundation (Cleveland, OH) using protocol number 1862. All mice used were on a C57BL/6NJ background, were maintained on a 14/10h light/dark cycle and had *ad libitum* access to food and water. Mice (housed 2-5/cage) did not have any prior history of drug administration, surgery, or behavioral testing.

### Primary human and murine astrocyte cultures

Primary human spinal cord astrocytes were obtained commercially from ScienCell Laboratories and grown according to provided protocols in complete ScienCell Astrocyte Medium. Briefly, astrocytes were isolated from the spinal cord and morphology was assessed by phase and relief contrast microscopy and GFAP positivity by immunofluorescence. Cell number, viability (≥ 70%), and proliferative potential (≥ 15 pd), and negative screening for potential biological contaminants was confirmed prior to cryopreservation. Following receipt, astrocytes were passaged and P3 cells were used for all studies. qPCR analysis for astrocyte purity was conducted prior to use in all experiments (**Supplemental Figure 1A**).

Primary murine spinal cord astrocytes were harvested as previously described^53^ with minor modifications. P2-P4 pups were euthanized and spinal cords harvested via hydro-ejection with PBS in ice cold, serum free DMEM containing 1% penicillin/streptomycin. The spinal cords were then digested with gentle agitation for 15 min in 0.25% trypsin-EDTA. Trypsinization was stopped by adding DMEM media supplemented with 10% heat inactivated FBS and 1% penicillin/streptomycin. Cells were then treated with 50μg/mL of DNase I for 3 min. Single cell suspensions were then made via trituration and cells were pelleted via centrifugation at 500x*g* for 5 min. Cells were resuspended in 10% DMEM and plated onto fibronectin-coated plates. Cells received a complete media change every other day until reaching 80% confluency. Astrocytes were mechanically separated from microglia and oligodendrocyte progenitor cells via orbital shaking.

### Post-mortem human MS tissue

Periventricular white matter from MS patients (**Table 1**) was collected according to the established rapid autopsy protocol approved by the Cleveland Clinic Institutional Review Board. MS patient tissue was prepared in 1cm thick slices using a guided box and was fixed in 4% paraformaldehyde (PFA) followed by sectioning for morphological and immunohistochemical analysis.

## Method details

### EAE Induction

Mice of mixed sex were induced for EAE at 9-10 weeks of age. C57BL/6NJ and *Batf2*^−/−^ mice were obtained commercially from The Jackson Laboratories and housed under specific pathogen-free conditions. *Batf2*^+/-^ mice were bred together to obtain littermate controls. C57BL/6NJ mice were bred together and used as controls. On day 0, mice were immunized s.c. with 100μg MOG_35-55_ emulsified in complete Freund’s adjuvant containing 400mg of heat killed Mycobacterium tuberculosis H37Ra using a standard emulsion (Hooke Laboratories). Pertussis toxin (100ng) (Hooke Laboratories) was injected i.p. on the day of immunization and two days later. Mice were monitored daily for clinical signs of disease as follows: 0, no observable signs; 1, limp tail; 2, limp tail and ataxia; 2.5, limp tail and knuckling of at least one limb; 3, paralysis of one limb; 3.5; partial paralysis of one limb and complete paralysis of the other; 4, complete hindlimb paralysis; 4.5, moribund; 5, death. Animals that did not develop clinical signs of EAE were excluded from the study. On day 25, mice were euthanized for histological studies via transcardial perfusion with PBS and 4% PFA. Mice were dissected to preserve spinal cord tissue integrity followed by fixation in 4% PFA overnight. Tissue was then transferred to a 30% sucrose solution for cryopreservation and spinal cords were frozen in O.C.T. Compound (Fisher Healthcare). Frozen, transverse sections (10μm) were slide mounted and stored at –80°C until immunofluorescent labeling.

### Immunofluorescent labeling

Tissue sections were blocked with 10% goat serum (Sigma) and 0.1% Triton X-100 for 1h at room temperature and then incubated with anti-MBP (Abcam), anti-GFAP (Invitrogen), CD45 PerCP-Cy5.5 (Biolegend), and anti-Caspase1 (Abclonal) primary antibodies overnight at 4°C. Tissue sections labeled with anti-IRF1 (Proteintech) were incubated in 1% Triton X-100 diluted in PBS for 20 min at room temperature prior to blocking and then incubated in primary antibody as previously described. Secondary antibodies conjugated to Alexa Fluor 488, Alex Fluor 555, and Alexa Fluor 647 (Invitrogen) were applied for 1h at room temperature as appropriate. Nuclei were counterstained with DAPI (Thermofisher) diluted in PBS. Sections were imaged using the 20x objective of a confocal microscope LSM 800 (Carl Zeiss). Representative images shown are illustrative of 3-6 images taken across two or more tissue sections at least 100μm apart per individual mouse. The mean positive area and Mander’s coefficient of colocalization were determined by setting thresholds using appropriate controls and quantified using ImageJ software (NIH). Lesion area was determined by tracing the area of mononuclear cell infiltration, using DAPI and CD45 staining, into the parenchyma using ImageJ software (NIH).

Free floating sections of human periventricular white matter were subjected to antigen retrieval by boiling the tissue briefly in 10μM citrate buffer. Sections were blocked with 5% donkey serum and 0.03% Triton X-100 (Sigma Aldrich) for 1h at room temperature. Demyelinated lesions were identified and characterized using immunostaining for MBP (Abcam) and Iba1 (Wako) as previously described^9,24^. Subsequent sections were exposed to antibodies specific for human BATF2 (Santa Cruz), IRF1 (Proteintech), Sox9 (R&D Systems), ALDH1L1 (Cell Signaling), and/or IBA1 (SPICA Dye 594-conjugated; Wako) for 4-5 days at 4°C. Sections were then washed with PBS-Triton-X-100, and secondary antibodies conjugated to Alexa Fluor 405, 488, and 647 (Thermofisher) were applied for 1h at room temperature as appropriate. Sections were then treated with 0.3% Sudan black in 70% ethanol for 3 min, imaged using the 10x, 20x, and 63x objectives of a confocal microscope LSM 800 (Carl Zeiss) and analyzed using ImageJ (NIH).

### RNA sequencing

Total RNA was collected from human primary spinal cord astrocytes (ScienCell) treated with recombinant human IFNγ, TNFα, IL-1β, IL-17, GMCSF, or media (Peprotech) for 24h using an RNeasy Mini Kit (Qiagen) according to the manufacturer’s instructions and quantified using a NanoDrop One spectrophotometer (Thermofisher). RNA was then sequenced at the Cleveland Clinic Lerner Research Institute Genomics Core. Count files were imported into R software and assessed for quality, normalized, and analyzed using an in-house pipeline to test for differential gene expression using the EdgeR Bionconductor library. Differential gene expression was determined for each gene using Cufflinks. PCA plots were generated using R software and the data set was further analyzed for Log2 fold change and plotted using Prism software.

### Quantitative PCR (qPCR)

Total RNA was collected from human or murine primary spinal cord astrocytes following media or cytokine treatment using an RNeasy Mini Kit (Qiagen) according to the manufacturer’s instructions and measured with a NanoDrop One spectrophotometer (Thermofisher). Total RNA was then normalized, treated with DNase I (Thermofisher), and reversed transcribed to cDNA using a TaqMan Reverse Transcription kit (Applied Biosciences). Prepared cDNA was then used as a template for qPCR using a QuantStudio 6 Real-Time PCR system (Thermofisher). Relative gene expression was quantified after normalizing to *GAPDH* expression. Primer sequences used are listed in the Key Resources Table.

### Chromatin immunoprecipitation (ChIP) sequencing and pathway analysis

Primary human astrocytes were stimulated with media or 10ng/mL IFNγ for 24h and fixed for 15 min in 1/10 volume Formaldehyde solution (11% Formaldehyde, 0.1M NaCl, 1mM EDTA pH 8.0, 50mM Hepes pH 7.9 in milliQ H_2_O) at room temperature. Fixation was stopped by adding 1/20 volume glycine solution (2.5M in milliQ H_2_O) for 5 min at room temperature. Cells were then scraped, transferred into 50mL conical tubes, and kept on ice. Tubes were centrifuged at 800x*g* at 4°C for 10 min. Pelleted cells were then resuspended in 10mL of chilled PBS-Igepal solution (0.5% Igepal in 1X PBS). Cells were repelleted as before and washed twice more with PBS-Igepal, adding 100uL of PMSF (1mM in ethanol) after the second wash to each sample. Pellets were then snap frozen on dry ice. ChIP sequencing and analysis was performed commercially by Active Motif Services. The resulting gene set associated with BATF2 chromatin binding was analyzed using IPA software (Ingenuity System Inc.) to examine canonical pathways regulated by BATF2 in media– and IFNγ-treated samples.

### Lysate preparation and Western blotting

Primary human spinal cord astrocytes were treated with media or 10ng/mL IFNγ for 24h. Cells were trypsonized with 0.05% Trypsin EDTA and homogenized in RIPA lysis buffer (50mM Tris-HCl pH=7.4, 150mM NaCl, 1mM EDTA, 1% Triton X-100, and 1:100 proteinase/phosphatase inhibitor). Samples were briefly triturated, vortexed for 3s, and incubated on ice for 30 min. For complete lysis, samples were frozen on dry ice for 3 min and thawed in wet ice for 15 min. A total of 3 freeze-thaw cycles were conducted. Samples were then centrifuged at 10,000xg for 30 min and supernatants were collected for immunoblotting. Protein levels were quantified using a Pierce BCA kit (Thermofisher) and normalized. 2x Laemmli sample buffer and β-Mercaptoethanol were added to normalized samples. 20μg of total protein lysate was resolved on Novex WedgeWell 4-20% Tris-Glycine gels and transferred to nitrocellulose membranes. The membranes were blocked with 5% low-fat powdered milk in Tris-buffered saline, 0.1% Tween® 20 (TBST) for 1h at room temperature and incubated overnight at 4°C with primary antibody. Membranes were blotted with anti-BATF2 (Thermofisher) and anti-β-actin (ThermoFisher) antibodies. The following day membranes were washed with TBST 3 times and incubated with HRP conjugated antibodies for 1h at room temperature. Membranes were then washed with TBST 3 times and imaged using the ChemiDoc MP imaging system (Bio-Rad) after activation with ECL substrate (Bio-Rad).

## Quantification and statistical analysis

### Statistical Analysis

Data and statistical analysis were performed using Prism 10.1.2 (Graphpad). Significance criteria includes: ns (not significant), *p*>0.05, \**p*<0.05, \*\**p*<0.01, \*\*\**p*<0.001, and \*\*\*\**p*<0.0001. Differences in means between greater than 2 groups were analyzed using a one-way ANOVA (with Tukey’s post hoc correction for multiple comparisons). For mixed groups, a two-way ANOVA (with Tukey’s post hoc correction for multiple comparisons) was used. EAE data were analyzed using the nonparametric Mann-Whitney *U* test. Graph bars indicate mean and error bars indicate standard error (SEM). For *in vivo* experiments, each *n* is representative of individual animals. For *in vitro* experiments, each *n* is representative of a separate, independent experiment. The *n* value for all experiments can be found in the provided Figure Legends.

---

**Supplemental Figure 1.**
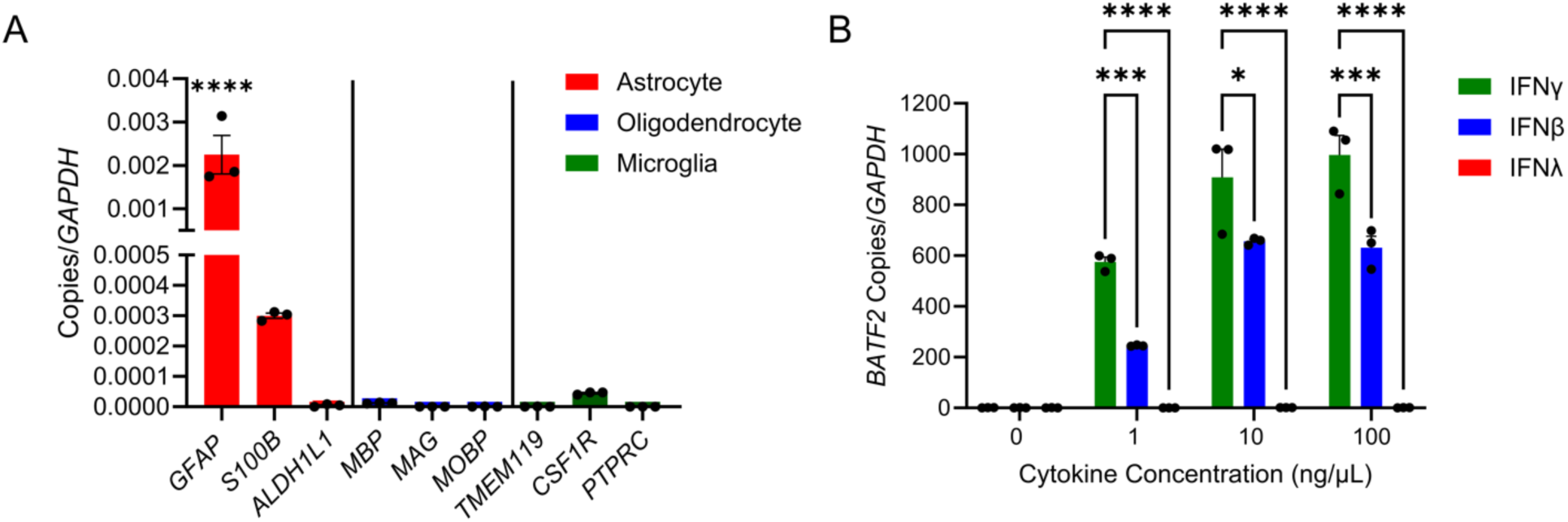
Validation of human spinal cord astrocytes and BATF2 interferon titration. (A) Transcript levels of astrocyte-(red), oligodendrocyte-(blue), and microglia-(green) specific genes expressed by untreated primary human spinal cord astrocytes. \*\*\*\**p* < 0.0001 by one-way ANOVA. Data points represent mean ± SEM (*n*=3). (B) *BATF2* transcript levels of human spinal cord astrocytes stimulated with 0ng/mL, 1ng/mL, 10ng/mL, or 100ng/mL of IFNγ, IFNϕ3, or IFNβ for 24h. \**p* < 0.05, \*\*\**p* < 0.001, \*\*\*\**p* < 0.0001 by two-way ANOVA. Data points represent mean ± SEM (*n*=3).

**Supplemental Figure 2.**
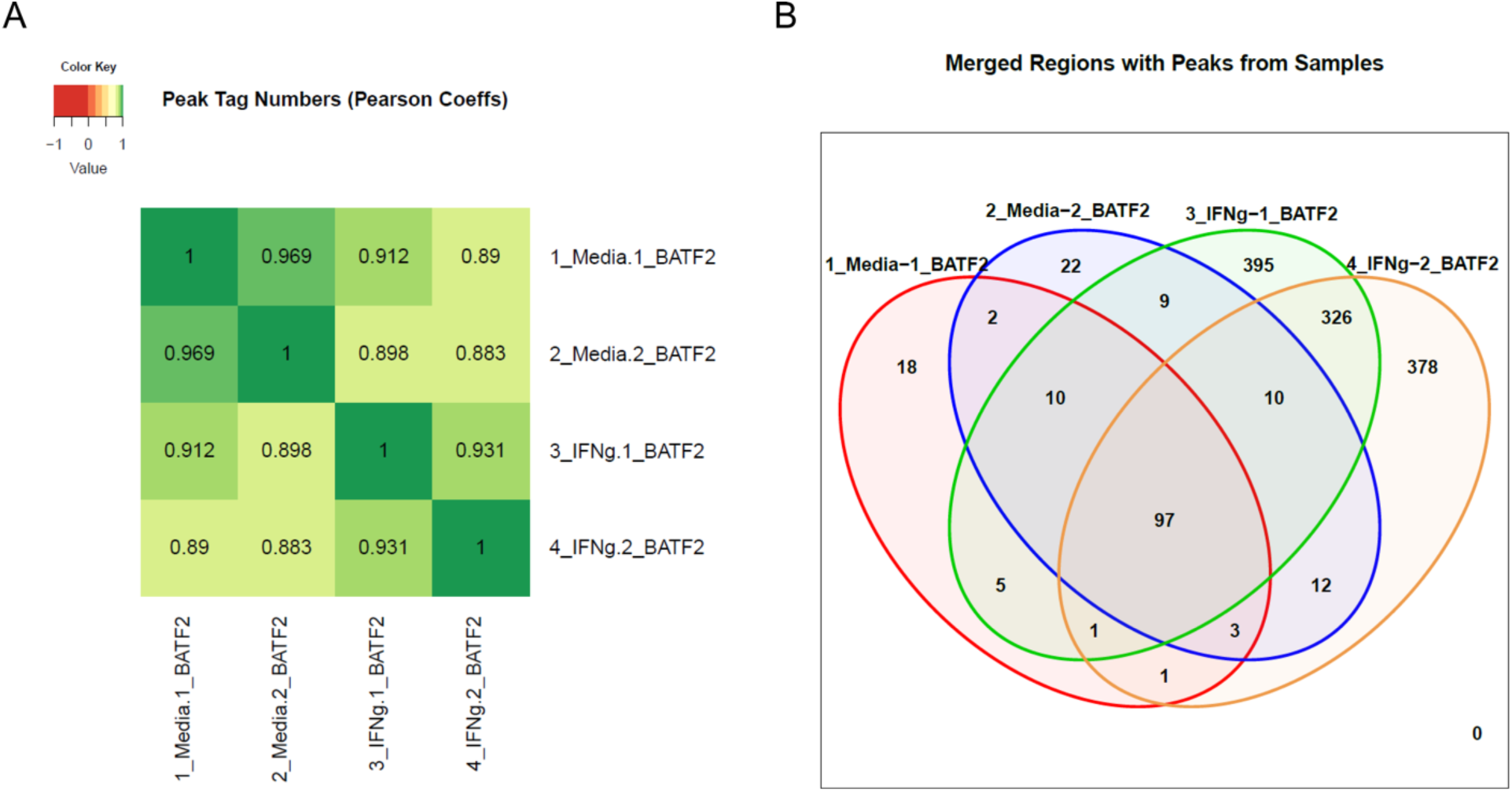
ChIP sequencing quality control and validation. (A) Pearson coefficient correlation plot of BATF2 peak tag numbers in primary human spinal cord astrocytes treated with 10ng/mL IFNγ or media for 24h. (B) Venn diagram of merged peak regions of human primary spinal cord astrocytes treated with 10ng/mL IFNγ or media for 24h. Data shown in A-E are representative of two independent samples per treatment.

**Supplemental Figure 3.**
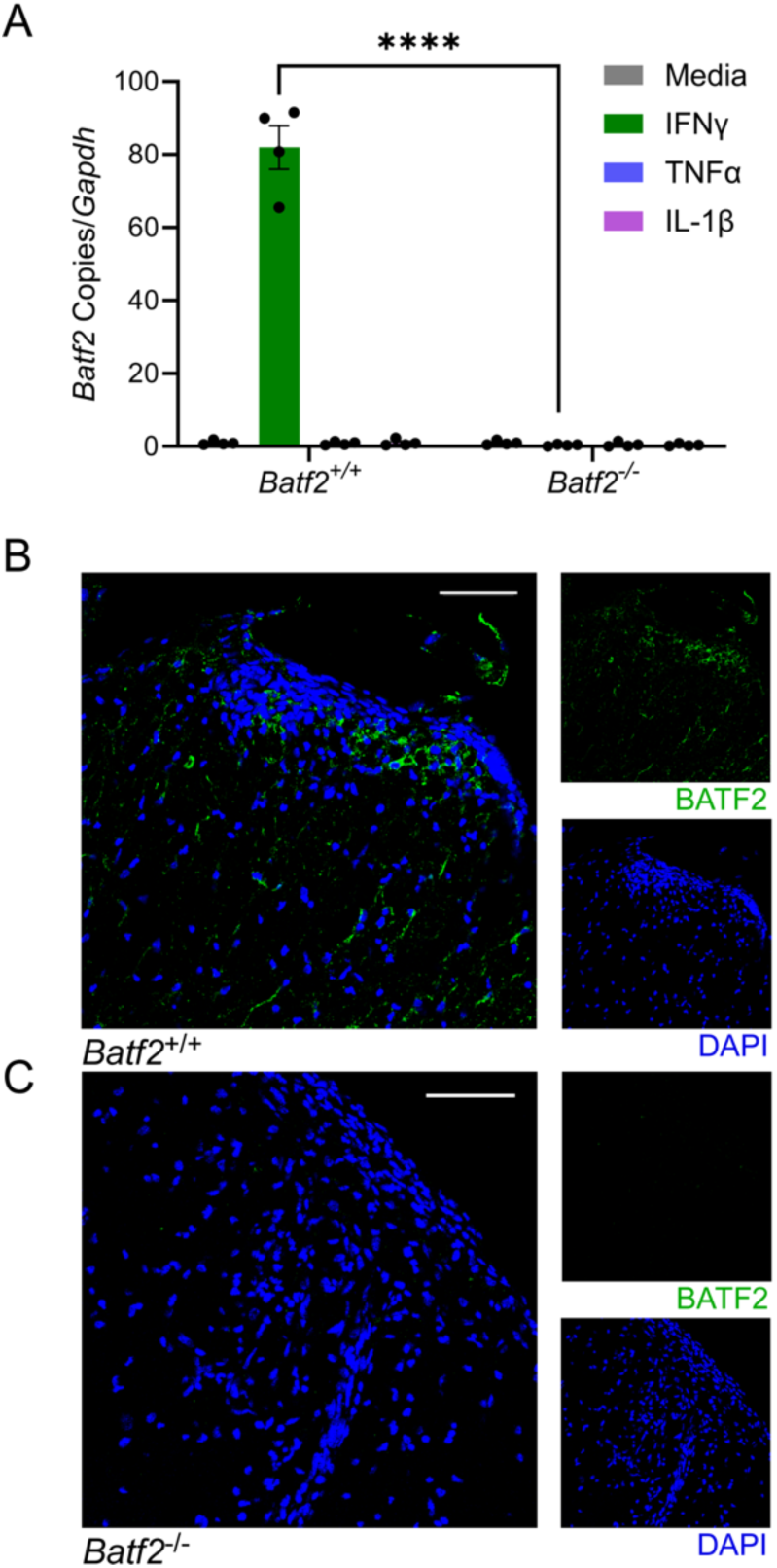
Validation of *Batf2*^−/−^ mouse. (A) Transcript levels of *Batf2* in *Batf2*^−/−^ and control murine spinal cord astrocytes treated with IFNγ or media for 48h. Individual data points were normalized to the media average and are representative of individual mice. \*\*\*\**p* < 0.0001 by two-way ANOVA comparing IFNγ treated samples. Data points represent mean ± SEM (*n*=4). (B-C) Spinal cord tissue from (B) *Batf2*^+/+^ and (C) *Batf2*^−/−^ mice labeled for BATF2 and nuclei counterstained with DAPI. Scale bars, 50μm.

**Supplemental Figure 4.**
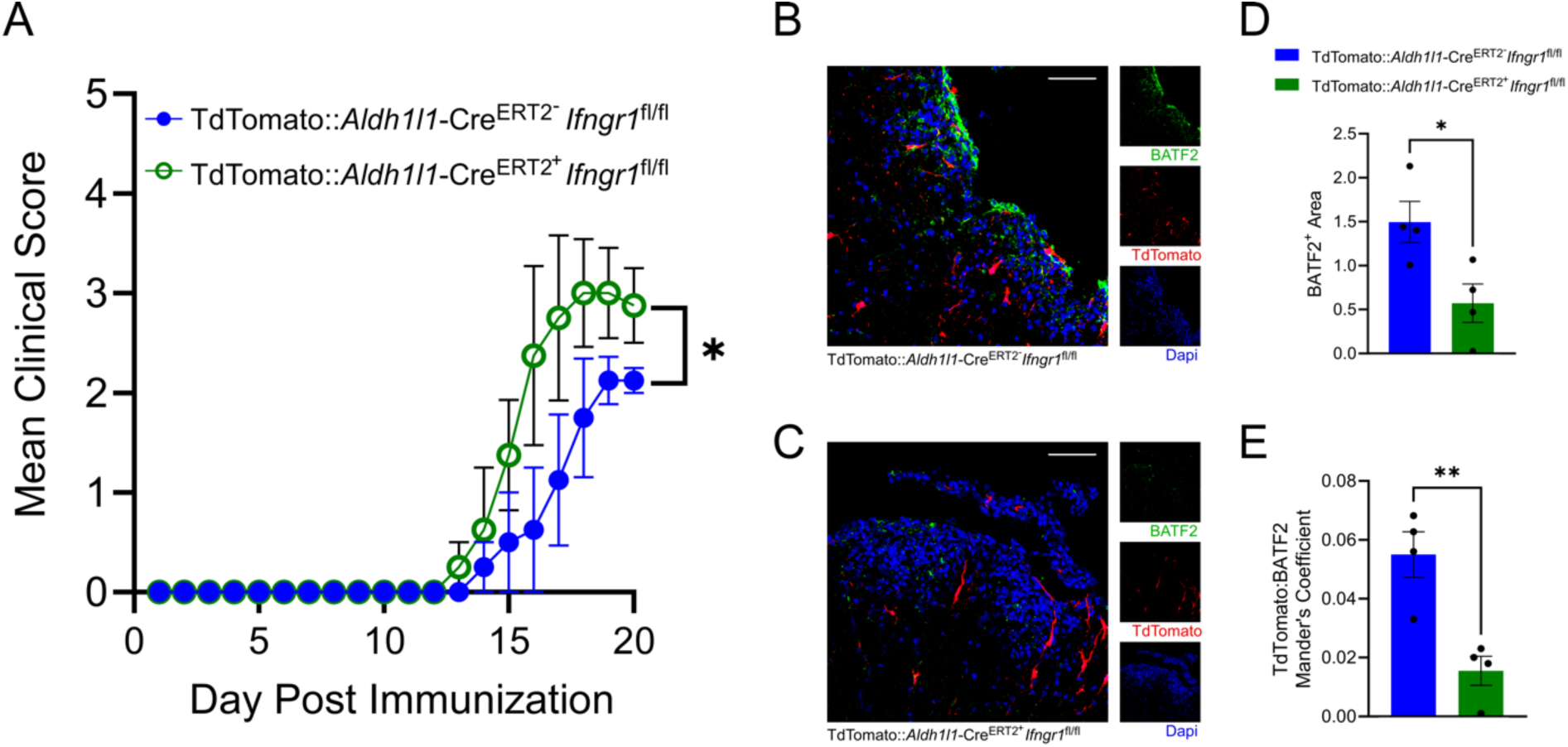
Astrocytic BATF2 expression is regulated by IFNγ *in vivo*. (A) Tamoxifen was injected into TdTomato::*Aldh1l1*-Cre^ERT2+^ *Ifngr1*^fl/fl^ and control mice two weeks prior to EAE induction. EAE clinical course was blindly monitored. Data was analyzed using the Mann–Whitney *U* test for nonparametric data. \**p* < 0.05 from day 14. Data points represent mean ± SEM (*n*=4 per genotype). 20 days post-immunization, mice were sacrificed and the CNS tissue was cryopreserved for immunofluorescent analysis. Ventral white matter tracts of the lumbar spinal cord were imaged using confocal microscopy. (B-C) Spinal cord tissue from (B) TdTomato::*Aldh1l1*-Cre^ERT2+^ *Ifngr1*^fl/fl^ and (C) control mice were labeled for BATF2 and nuclei counterstained with DAPI. Scale bars, 50μm. (D-E) Quantification of BATF2 area and colocalization between BATF2 and TdTomato in TdTomato::*Aldh1l1*-Cre^ERT2+^ *Ifngr1*^fl/fl^ (green) and control (blue) mice. \*\**p* < 0.01, \*\*\**p* < 0.001 by a two-tailed Student’s *t* test for each set of bars. Bars represent mean ± SEM (*n*=4 per genotype).

## References

1. Escartin, C., Galea, E., Lakatos, A., O’Callaghan, J.P., Petzold, G.C., Serrano-Pozo, A., Steinhauser, C., Volterra, A., Carmignoto, G., Agarwal, A., et al. (2021). Reactive astrocyte nomenclature, definitions, and future directions. Nat Neurosci 24, 312–325. 10.1038/s41593-020-00783-4.

2. Brosnan, C.F., and Raine, C.S. (2013). The astrocyte in multiple sclerosis revisited. Glia 61, 453–465. 10.1002/glia.22443.

3. Olsson, T. (1992). Cytokines in neuroinflammatory disease: role of myelin autoreactive T cell production of interferon-gamma. Journal of neuroimmunology 40, 211–218.

4. Fletcher, J.M., Lalor, S.J., Sweeney, C.M., Tubridy, N., and Mills, K.H. (2010). T cells in multiple sclerosis and experimental autoimmune encephalomyelitis. Clin Exp Immunol 162, 1–11. 10.1111/j.1365-2249.2010.04143.x.

5. Hashioka, S., McGeer, E.G., Miyaoka, T., Wake, R., Horiguchi, J., and McGeer, P.L. (2015). Interferon-γ-induced neurotoxicity of human astrocytes. CNS Neurol Disord Drug Targets 14, 251–256. 10.2174/1871527314666150217122305.

6. Sun, L., Li, Y., Jia, X., Wang, Q., Li, Y., Hu, M., Tian, L., Yang, J., Xing, W., Zhang, W., et al. (2017). Neuroprotection by IFN-γ via astrocyte-secreted IL-6 in acute neuroinflammation. Oncotarget 8, 40065–40078. 10.18632/oncotarget.16990.

7. Hindinger, C., Bergmann, C.C., Hinton, D.R., Phares, T.W., Parra, G.I., Hussain, S., Savarin, C., Atkinson, R.D., and Stohlman, S.A. (2012). IFN-γ signaling to astrocytes protects from autoimmune mediated neurological disability. PLoS One 7, e42088. 10.1371/journal.pone.0042088.

8. Savarin, C., Hinton, D.R., Valentin-Torres, A., Chen, Z., Trapp, B.D., Bergmann, C.C., and Stohlman, S.A. (2015). Astrocyte response to IFN-γ limits IL-6-mediated microglia activation and progressive autoimmune encephalomyelitis. J Neuroinflammation 12, 79. 10.1186/s12974-015-0293-9.

9. Smith, B.C., Tinkey, R.A., Brock, O.D., Mariam, A., Habean, M.L., Dutta, R., and Williams, J.L. (2023). Astrocyte interferon-gamma signaling dampens inflammation during chronic central nervous system autoimmunity via PD-L1. Journal of neuroinflammation 20, 234. 10.1186/s12974-023-02917-4.

10. Smith, B.C., Sinyuk, M., Jenkins, J.E., 3rd, Psenicka, M.W., and Williams, J.L. (2020). The impact of regional astrocyte interferon-gamma signaling during chronic autoimmunity: a novel role for the immunoproteasome. Journal of neuroinflammation 17, 184. 10.1186/s12974-020-01861-x.

11. Linnerbauer, M., Beyer, T., Nirschl, L., Farrenkopf, D., Losslein, L., Vandrey, O., Peter, A., Tsaktanis, T., Kebir, H., Laplaud, D., et al. (2023). PD-L1 positive astrocytes attenuate inflammatory functions of PD-1 positive microglia in models of autoimmune neuroinflammation. Nat Commun 14, 5555. 10.1038/s41467-023-40982-8.

12. Sanmarco, L.M., Wheeler, M.A., Gutierrez-Vazquez, C., Polonio, C.M., Linnerbauer, M., Pinho-Ribeiro, F.A., Li, Z., Giovannoni, F., Batterman, K.V., Scalisi, G., et al. (2021). Gut-licensed IFNgamma(+) NK cells drive LAMP1(+)TRAIL(+) anti-inflammatory astrocytes. Nature 590, 473–479. 10.1038/s41586-020-03116-4.

13. Su, Z.Z., Lee, S.G., Emdad, L., Lebdeva, I.V., Gupta, P., Valerie, K., Sarkar, D., and Fisher, P.B. (2008). Cloning and characterization of SARI (suppressor of AP-1, regulated by IFN). Proceedings of the National Academy of Sciences of the United States of America 105, 20906–20911. 10.1073/pnas.0807975106.

14. Kitada, S., Kayama, H., Okuzaki, D., Koga, R., Kobayashi, M., Arima, Y., Kumanogoh, A., Murakami, M., Ikawa, M., and Takeda, K. (2017). BATF2 inhibits immunopathological Th17 responses by suppressing Il23a expression during Trypanosoma cruzi infection. J Exp Med 214, 1313–1331. 10.1084/jem.20161076.

15. Dai, L., Liu, Y., Cheng, L., Wang, H., Lin, Y., Shi, G., Dong, Z., Li, J., Fan, P., Wang, Q., et al. (2019). SARI attenuates colon inflammation by promoting STAT1 degradation in intestinal epithelial cells. Mucosal Immunol 12, 1130–1140. 10.1038/s41385-019-0178-9.

16. Roy, S., Guler, R., Parihar, S.P., Schmeier, S., Kaczkowski, B., Nishimura, H., Shin, J.W., Negishi, Y., Ozturk, M., Hurdayal, R., et al. (2015). Batf2/Irf1 induces inflammatory responses in classically activated macrophages, lipopolysaccharides, and mycobacterial infection. J Immunol 194, 6035–6044. 10.4049/jimmunol.1402521.

17. van der Geest, R., Peñaloza, H.F., Xiong, Z., Gonzalez-Ferrer, S., An, X., Li, H., Fan, H., Tabary, M., Nouraie, S.M., Zhao, Y., et al. (2023). BATF2 enhances proinflammatory cytokine responses in macrophages and improves early host defense against pulmonary Klebsiella pneumoniae infection. Am J Physiol Lung Cell Mol Physiol 325, L604–l616. 10.1152/ajplung.00441.2022.

18. Guler, R., Mpotje, T., Ozturk, M., Nono, J.K., Parihar, S.P., Chia, J.E., Abdel Aziz, N., Hlaka, L., Kumar, S., Roy, S., et al. (2019). Batf2 differentially regulates tissue immunopathology in Type 1 and Type 2 diseases. Mucosal Immunol 12, 390–402. 10.1038/s41385-018-0108-2.

19. Furlan, R., Martino, G., Galbiati, F., Poliani, P.L., Smiroldo, S., Bergami, A., Desina, G., Comi, G., Flavell, R., Su, M.S., and Adorini, L. (1999). Caspase-1 regulates the inflammatory process leading to autoimmune demyelination. J Immunol 163, 2403–2409.

20. Ren, Z., Wang, Y., Liebenson, D., Liggett, T., Goswami, R., Stefoski, D., and Balabanov, R. (2011). IRF-1 signaling in central nervous system glial cells regulates inflammatory demyelination. J Neuroimmunol 233, 147–159. 10.1016/j.jneuroim.2011.01.001.

21. Ren, Z., Wang, Y., Tao, D., Liebenson, D., Liggett, T., Goswami, R., Clarke, R., Stefoski, D., and Balabanov, R. (2011). Overexpression of the dominant-negative form of interferon regulatory factor 1 in oligodendrocytes protects against experimental autoimmune encephalomyelitis. J Neurosci 31, 8329–8341. 10.1523/jneurosci.1028-11.2011.

22. Buch, T., Uthoff-Hachenberg, C., and Waisman, A. (2003). Protection from autoimmune brain inflammation in mice lacking IFN-regulatory factor-1 is associated with Th2-type cytokines. Int Immunol 15, 855–859. 10.1093/intimm/dxg086.

23. Tada, Y., Ho, A., Matsuyama, T., and Mak, T.W. (1997). Reduced incidence and severity of antigen-induced autoimmune diseases in mice lacking interferon regulatory factor-1. J Exp Med 185, 231–238. 10.1084/jem.185.2.231.

24. Smith, B.C., Sinyuk, M., Jenkins, J.E., 3rd, Psenicka, M.W., and Williams, J.L. (2020). The impact of regional astrocyte interferon-γ signaling during chronic autoimmunity: a novel role for the immunoproteasome. J Neuroinflammation 17, 184. 10.1186/s12974-020-01861-x.

25. Lycklama, G., Thompson, A., Filippi, M., Miller, D., Polman, C., Fazekas, F., and Barkhof, F. (2003). Spinal-cord MRI in multiple sclerosis. The Lancet. Neurology 2, 555–562.

26. Nijeholt, G.J., van Walderveen, M.A., Castelijns, J.A., van Waesberghe, J.H., Polman, C., Scheltens, P., Rosier, P.F., Jongen, P.J., and Barkhof, F. (1998). Brain and spinal cord abnormalities in multiple sclerosis. Correlation between MRI parameters, clinical subtypes and symptoms. Brain: a journal of neurology 121 ( *Pt 4*), 687–697.

27. Handunnetthi, L., Ramagopalan, S.V., Ebers, G.C., and Knight, J.C. (2010). Regulation of major histocompatibility complex class II gene expression, genetic variation and disease. Genes Immun 11, 99–112. 10.1038/gene.2009.83.

28. Sugiyama, M., Kikuchi, A., Misu, H., Igawa, H., Ashihara, M., Kushima, Y., Honda, K., Suzuki, Y., Kawabe, Y., Kaneko, S., and Takamura, T. (2018). Inhibin βE (INHBE) is a possible insulin resistance-associated hepatokine identified by comprehensive gene expression analysis in human liver biopsy samples. PLoS One 13, e0194798. 10.1371/journal.pone.0194798.

29. Dinarello, C.A., Novick, D., Kim, S., and Kaplanski, G. (2013). Interleukin-18 and IL-18 binding protein. Front Immunol 4, 289. 10.3389/fimmu.2013.00289.

30. Kayama, H., Tani, H., Kitada, S., Opasawatchai, A., Okumura, R., Motooka, D., Nakamura, S., and Takeda, K. (2019). BATF2 prevents T-cell-mediated intestinal inflammation through regulation of the IL-23/IL-17 pathway. Int Immunol 31, 371–383. 10.1093/intimm/dxz014.

31. Oldfield, A.J., Yang, P., Conway, A.E., Cinghu, S., Freudenberg, J.M., Yellaboina, S., and Jothi, R. (2014). Histone-fold domain protein NF-Y promotes chromatin accessibility for cell type-specific master transcription factors. Mol Cell 55, 708–722. 10.1016/j.molcel.2014.07.005.

32. Oldfield, A.J., Henriques, T., Kumar, D., Burkholder, A.B., Cinghu, S., Paulet, D., Bennett, B.D., Yang, P., Scruggs, B.S., Lavender, C.A., et al. (2019). NF-Y controls fidelity of transcription initiation at gene promoters through maintenance of the nucleosome-depleted region. Nat Commun 10, 3072. 10.1038/s41467-019-10905-7.

33. Fleming, J.D., Pavesi, G., Benatti, P., Imbriano, C., Mantovani, R., and Struhl, K. (2013). NF-Y coassociates with FOS at promoters, enhancers, repetitive elements, and inactive chromatin regions, and is stereo-positioned with growth-controlling transcription factors. Genome Res 23, 1195–1209. 10.1101/gr.148080.112.

34. Wei, C.J., Li, Y.L., Zhu, Z.L., Jia, D.M., Fan, M.L., Li, T., Wang, X.J., Li, Z.G., and Ma, H.S. (2019). Inhibition of activator protein 1 attenuates neuroinflammation and brain injury after experimental intracerebral hemorrhage. CNS Neurosci Ther 25, 1182–1188. 10.1111/cns.13206.

35. Yamazaki, S., Tanaka, Y., Araki, H., Kohda, A., Sanematsu, F., Arasaki, T., Duan, X., Miura, F., Katagiri, T., Shindo, R., et al. (2017). The AP-1 transcription factor JunB is required for Th17 cell differentiation. Sci Rep 7, 17402. 10.1038/s41598-017-17597-3.

36. Bagnoud, M., Briner, M., Remlinger, J., Meli, I., Schuetz, S., Pistor, M., Salmen, A., Chan, A., and Hoepner, R. (2020). c-Jun N-Terminal Kinase as a Therapeutic Target in Experimental Autoimmune Encephalomyelitis. Cells 9. 10.3390/cells9102154.

37. Groves, A., Kihara, Y., Jonnalagadda, D., Rivera, R., Kennedy, G., Mayford, M., and Chun, J. (2018). A Functionally Defined In Vivo Astrocyte Population Identified by c-Fos Activation in a Mouse Model of Multiple Sclerosis Modulated by S1P Signaling: Immediate-Early Astrocytes (ieAstrocytes). eNeuro 5. 10.1523/eneuro.0239-18.2018.

38. Schraml, B.U., Hildner, K., Ise, W., Lee, W.L., Smith, W.A., Solomon, B., Sahota, G., Sim, J., Mukasa, R., Cemerski, S., et al. (2009). The AP-1 transcription factor Batf controls T(H)17 differentiation. Nature 460, 405–409. 10.1038/nature08114.

39. Karwacz, K., Miraldi, E.R., Pokrovskii, M., Madi, A., Yosef, N., Wortman, I., Chen, X., Watters, A., Carriero, N., Awasthi, A., et al. (2017). Critical role of IRF1 and BATF in forming chromatin landscape during type 1 regulatory cell differentiation. Nature immunology 18, 412–421. 10.1038/ni.3683.

40. Li, W., Viengkhou, B., Denyer, G., West, P.K., Campbell, I.L., and Hofer, M.J. (2018). Microglia have a more extensive and divergent response to interferon-α compared with astrocytes. Glia 66, 2058–2078. 10.1002/glia.23460.

41. Fortunato, G., Calcagno, G., Bresciamorra, V., Salvatore, E., Filla, A., Capone, S., Liguori, R., Borelli, S., Gentile, I., Borrelli, F., et al. (2008). Multiple sclerosis and hepatitis C virus infection are associated with single nucleotide polymorphisms in interferon pathway genes. J Interferon Cytokine Res 28, 141–152. 10.1089/jir.2007.0049.

42. Liddelow, S.A., Guttenplan, K.A., Clarke, L.E., Bennett, F.C., Bohlen, C.J., Schirmer, L., Bennett, M.L., Münch, A.E., Chung, W.S., Peterson, T.C., et al. (2017). Neurotoxic reactive astrocytes are induced by activated microglia. Nature 541, 481–487. 10.1038/nature21029.

43. Wang, H., Song, G., Chuang, H., Chiu, C., Abdelmaksoud, A., Ye, Y., and Zhao, L. (2018). Portrait of glial scar in neurological diseases. Int J Immunopathol Pharmacol 31, 2058738418801406. 10.1177/2058738418801406.

44. Bush, T.G., Puvanachandra, N., Horner, C.H., Polito, A., Ostenfeld, T., Svendsen, C.N., Mucke, L., Johnson, M.H., and Sofroniew, M.V. (1999). Leukocyte infiltration, neuronal degeneration, and neurite outgrowth after ablation of scar-forming, reactive astrocytes in adult transgenic mice. Neuron 23, 297–308.

45. Faulkner, J.R., Herrmann, J.E., Woo, M.J., Tansey, K.E., Doan, N.B., and Sofroniew, M.V. (2004). Reactive astrocytes protect tissue and preserve function after spinal cord injury. J Neurosci 24, 2143–2155. 10.1523/JNEUROSCI.3547-03.2004.

46. Toft-Hansen, H., Füchtbauer, L., and Owens, T. (2011). Inhibition of reactive astrocytosis in established experimental autoimmune encephalomyelitis favors infiltration by myeloid cells over T cells and enhances severity of disease. Glia 59, 166–176. 10.1002/glia.21088.

47. Liu, Z., Li, Y., Cui, Y., Roberts, C., Lu, M., Wilhelmsson, U., Pekny, M., and Chopp, M. (2014). Beneficial effects of gfap/vimentin reactive astrocytes for axonal remodeling and motor behavioral recovery in mice after stroke. Glia 62, 2022–2033. 10.1002/glia.22723.

48. Zsurka, G., Appel, M.L.T., Nastaly, M., Hallmann, K., Hansen, N., Nass, D., Baumgartner, T., Surges, R., Hartmann, G., Bartok, E., and Kunz, W.S. (2023). Loss of the Immunomodulatory Transcription Factor BATF2 in Humans Is Associated with a Neurological Phenotype. Cells 12. 10.3390/cells12020227.

49. Ho, H.H., Antoniv, T.T., Ji, J.D., and Ivashkiv, L.B. (2008). Lipopolysaccharide-induced expression of matrix metalloproteinases in human monocytes is suppressed by IFN-gamma via superinduction of ATF-3 and suppression of AP-1. J Immunol 181, 5089–5097. 10.4049/jimmunol.181.7.5089.

50. Hu, X., Paik, P.K., Chen, J., Yarilina, A., Kockeritz, L., Lu, T.T., Woodgett, J.R., and Ivashkiv, L.B. (2006). IFN-gamma suppresses IL-10 production and synergizes with TLR2 by regulating GSK3 and CREB/AP-1 proteins. Immunity 24, 563–574. 10.1016/j.immuni.2006.02.014.

51. Radzioch, D., and Varesio, L. (1991). c-fos mRNA expression in macrophages is downregulated by interferon-gamma at the posttranscriptional level. Mol Cell Biol 11, 2718–2722. 10.1128/mcb.11.5.2718-2722.1991.

52. Hu, X., Chen, J., Wang, L., and Ivashkiv, L.B. (2007). Crosstalk among Jak-STAT, Toll-like receptor, and ITAM-dependent pathways in macrophage activation. J Leukoc Biol 82, 237–243. 10.1189/jlb.1206763.

53. Sinyuk, M., and Williams, J.L. (2020). Dissection and Isolation of Murine Glia from Multiple Central Nervous System Regions. Journal of visualized experiments: JoVE. 10.3791/61345.

